# Comparative transcriptomics of seven *Impatiens* species

**DOI:** 10.1101/2021.09.16.460688

**Authors:** Mária Šurinová, Štepán Stočes, Tomáš Dostálek, Andrea Jarošová, Zuzana Münzbergová

## Abstract

*Impatiens* is a genus containing more than 1000 species. Thanks to its size, it is a unique system for studying species diversification in natural populations. This study focused on the characterization of novel transcriptomes from seven *Impatiens* species originating from Nepal. Leave transcriptome of *Impatines balsamina* L.*, I. racemosa* DC.*, I. bicornuta* Wall*, I. falcifer* Hook*, I. devendrae* Pusalkar, *I. scullyi* Hook and *I. scabrida* DC were sequenced and compared. Reads were *de novo* assembled and aligned to 92 035-226 081 contigs. We identified 14 728 orthology groups shared among all the species and 3 020 which were unique to a single species. In single species, 2536-3009 orthology groups were under selection from which 767 were common for all species. Six of the seven investigated species shared 77% of gene families with *I. bicornuta* being the most distinct species. Specific gene families involved in response to different environmental stimuli were closely described. *Impatiens bicornuta* selection profile shared selection on zing finger protein structures and flowering regulation and stress response proteins with the other investigated species. Overall, the study showed substantial similarity in patterns of selections on transcribed genes across the species suggesting similar evolutionary pressures. This suggests that the species group may have evolved via adaptive radiation.

## Introduction

Most plant genera contain small number of species with only 57 genera (out of total of more than 13 000) of flowering plants containing more than 500 species (1). Two main mechanisms have been proposed to explain the high diversity of the species rich genera. The first is rapid radiation, which typically occurs in islands or mountains (2). It is described as a process of rapid diversification of an ancestral species via occupation of niches in newly available ecological space (3). The diversification is relatively fast and results in development of new forms adapted to new environments (2). The second proposed process is diversity accumulation in evolution, which in comparison to rapid radiation, is a slower process (4). At the genome level, species differentiation is often linked to polyploidization, genome rearrangement, recombination and complementing mutations (5). These mechanisms are studied for a long time (6–9). On the other hand, studies exploring plant species diversification from the perspective of gene expression are still rare (but see (10). The recent increase in the number of such studies occurred thanks to rise of next-generation sequencing methods allowing to expand whole transcriptome studies to natural, non-model organisms.

From a molecular perspective, species rich genera have been studied very poorly on DNA level and even less on RNA level. Most of the genetic studies comparing multiple species within larger genera are performed to understand species phylogeny by sequencing few genomic regions (11–13). These regions are inadequate to help explaining change in genetic mechanisms leading to adaptation. Knowledge of whole genome or transcriptome sequences is crucial. Gene annotation allows to identify gene functions, regulatory and biochemical pathways for known but also for newly identified genes. Genes are coding information behind the biological networks and this information brings essential insights into protein regulatory interactions that determine biochemical and physiological features of a cell, a tissue and an individual (14).

Comparative transcriptomes are mainly available for agricultural species on intraspecies level (15–18). Studies comparing transcriptomes of multiple natural species are rare (19, 20). Using RNA sequencing to compare different species has several advantages compared to whole genome sequencing. It allows to study only a regulatory part of the genome allowing to focus on the functional differences among the species and reduce the overall sequencing costs.

Comparative studies of natural populations are needed to better elucidate modulation of the genome to understand which mechanisms were involved in species differentiation.

Genus *Impatiens* L. (Balsaminaceae) consisting of >1000 species is a good example of a large genus (21). The genus is distributed mainly in tropical regions of the Old World and subtropics (22, 23). This genus is widespread from sea level to 4000 m above sea level and several species of the genus have become invasive in different parts of the world (24).

Because of its large distribution and high diversity, evolution and adaptation of this genus is widely studied from ecological perspective (25–33). On the other hand, genus *Impatiens* is not deeply studied at the molecular level. No whole-genome sequencing data are available, only six chloroplast genomes have been sequenced so far (34–39) and recently short data report for leaf transcriptome of *Impatience balsamina* was published (40). A combination of broad ecological and genetic research has unique potential to elucidate the importance of environmental factors in species differentiation processes.

In this study, we present the first comprehensive analysis of leaf transcriptomes of seven *Impatiens* species. In all seven species, RNA from leaf tissue of select individuals from an altitudinal gradient in Nepal was sequenced using Illumina short reads and assembled *de novo* into transcriptomes and annotated. A comparative study followed to characterize the gene content and identify patterns of selection among orthologous gene families to understand the genetic contribution to the evolution of the genus. Specifically, we asked 1) Do the species differ in transcriptome size and in transcriptome profiles in the de novo sequenced transcriptomes? 2) Do the identified genes under positive selection differ among species?

## Material and Methods

### Studied species

For our comparative transcriptome study, we used seven *Impatiens* species growing along altitudinal gradient in Nepal, Himalayas: *Impatines balsamina* L.*, I. racemosa* DC. *, I. bicornuta* Wall*, I. falcifer* Hook*, Impatiens aff. I. devendrae* Pusalkar (later referred as *I. devendrae*), *I. scullyi* Hook and *I. scabrida* DC. In recent revision, Akiyama and Ohba (41) suggested that name *I. tricornis* Lindl. should be used instead of *I. scabrida*. In this paper, we are using *I. scabrida* name because of continuity with previous studies. Species of genus *Impatiens* prefer wetter places having low tolerance to long term drought or long exposure to direct sunlight (42). We selected species differing in their altitudinal distribution in order to present species with different temperature niches (31) to investigate possible corresponding changes in transcriptomes. The selection of species was partly limited by the ability of the plants to germinate and survive in all our experimental conditions (see below).

Phylogenetic relationships among the studied species including other members of the genus *Impatiens* is well studied (23,43,44). Published phylogeny (43) defining phylogenetic structure based on sequence and morphological data, identified seven subclades (A-G) within the genus. All species studied here except *I. balsamina* belong to subclade B, *I. balsamina* belongs to subclade G.

### Plant material

Seeds of model species were collected from natural populations in autumn 2017 in Nepal (Supplementary Table S1) by M. Rokaya for purpose of our previous studies (26). Permission for the collection of plant material was obtained from the Department of National Parks and Wildlife Conservation in Nepal to Dr. Rokaya. Dr. Rokaya identified collected individuals in collaboration with Dr. Wojciech Adamowski (University of Warsaw). Voucher specimens are available in the Herbarium of the Institute of Botany of ASCR: PRA-20484, PRA-20486, PRA-20488, PRA-20490, PRA-20492, PRA-20494, PRA-20496. Seeds of each species were collected from at least five plants from one population consisting of at least several tens of individuals.

**Table 1.**
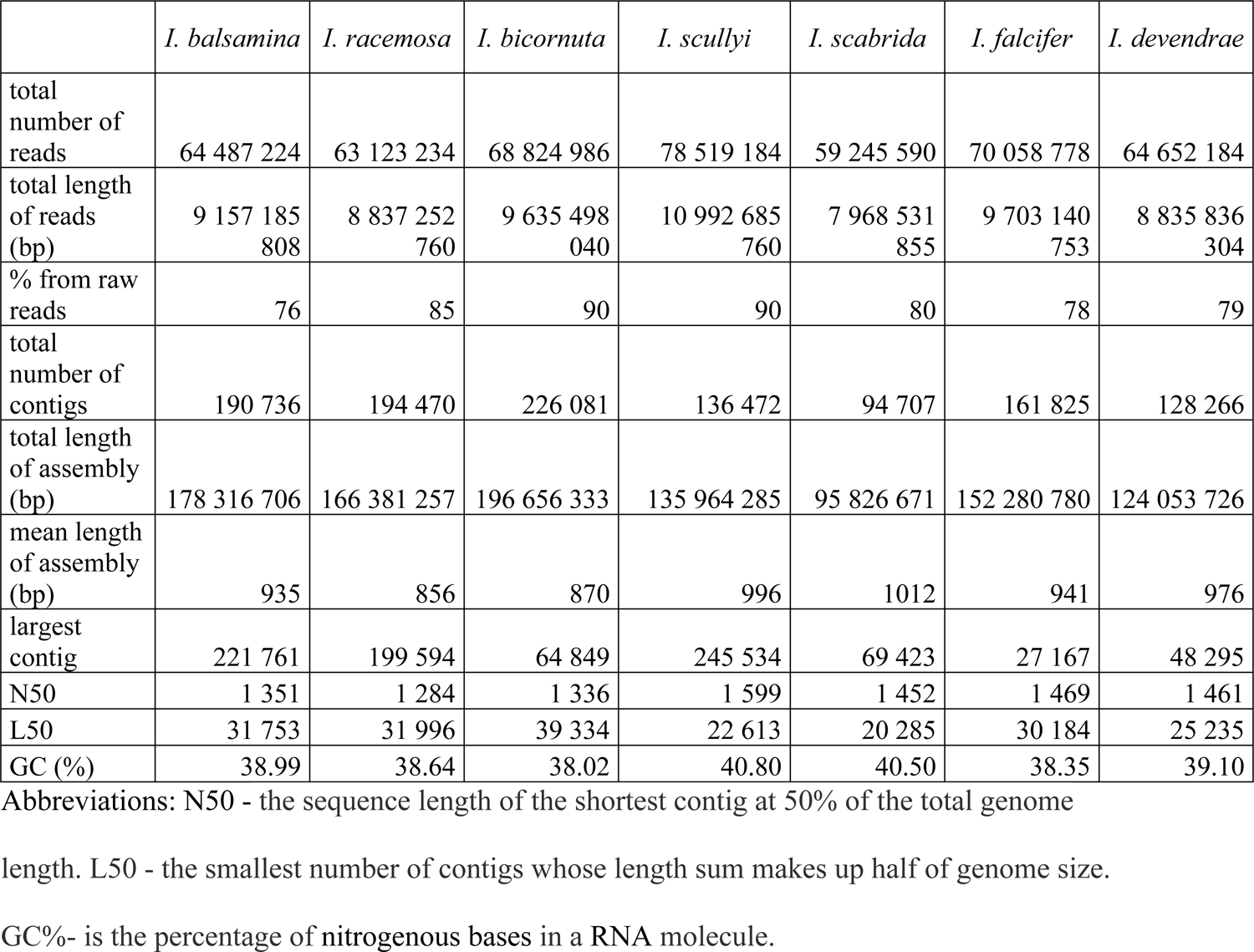
Summary statistics of sequencing data and de novo transcriptome assembly of seven *Impatiens* species.

|  | *I. balsamina* | *I. racemosa* | *I. bicornuta* | *I. scullyi* | *I. scabrida* | *I. falcifer* | *I. devendrae* |
| --- | --- | --- | --- | --- | --- | --- | --- |
| total number of reads | 64 487 224 | 63 123 234 | 68 824 986 | 78 519 184 | 59 245 590 | 70 058 778 | 64 652 184 |
| total length of reads (bp) | 9 157 185 808 | 8 837 252 760 | 9 635 498 040 | 10 992 685 760 | 7 968 531 855 | 9 703 140 753 | 8 835 836 304 |
| % from raw reads | 76 | 85 | 90 | 90 | 80 | 78 | 79 |
| total number of contigs | 190 736 | 194 470 | 226 081 | 136 472 | 94 707 | 161 825 | 128 266 |
| total length of assembly (bp) | 178 316 706 | 166 381 257 | 196 656 333 | 135 964 285 | 95 826 671 | 152 280 780 | 124 053 726 |
| mean length of assembly (bp) | 935 | 856 | 870 | 996 | 1012 | 941 | 976 |
| largest contig | 221 761 | 199 594 | 64 849 | 245 534 | 69 423 | 27 167 | 48 295 |
| N50 | 1 351 | 1 284 | 1 336 | 1 599 | 1 452 | 1 469 | 1 461 |
| L50 | 31 753 | 31 996 | 39 334 | 22 613 | 20 285 | 30 184 | 25 235 |
| GC (%) | 38.99 | 38.64 | 38.02 | 40.80 | 40.50 | 38.35 | 39.10 |

Abbreviations: N50 - the sequence length of the shortest contig at 50% of the total genome length. L50 - the smallest number of contigs whose length sum makes up half of genome size. GC%- is the percentage of nitrogenous bases in a RNA molecule.

We cultivated the plants in the growth chambers set to the conditions which plants are experiencing in their natural range. Two extremes were selected: (1) cold regime (mean, minimum and maximum temperatures of 12, 6 and 17.5 °C) corresponding to temperatures from March to June at 2,700 m a. s. l., representing the median of the higher altitudinal range of *Impatiens* species in Nepal, and (2) warm regime (mean, minimum and maximum temperatures of 21, 15 and 25 °C) corresponding to temperatures from March to June at 1,800 m a. s. l., representing the median of the lower altitudinal range of *Impatiens* species in Nepal in 2050 as predicted by the global climate model MIRO5C under the greenhouse gas concentration trajectory RCP8.5. The growth chambers were set to 12-h/12-h light/dark cycle, 250 µmol·m-2·s-1 light intensity. Temperature data were obtained from WorldClim database. We used mean temperatures from March to June since it represents premonsoon period when most *Impatiens* species germinate and start to grow. For further details on plant cultivation see (26, 31) using the same plants for studies dealing with species trait variation and attractivity to herbivores.

Fourteen samples were collected for RNA extractions (seven investigated species in two regimes). Fully expanded leaves from adult individuals were collected at the stage of flower buds. For each species and temperature regime, the exact age of plants and the time of sampling is different, but all the plants are in the same phenological stage. Leaves were collected during the light period in the same part of the day and immediately frozen in liquid nitrogen.

### RNA sequencing

RNA was extracted according to (45) with the following changes. 2.0 M sodium chloride and 25 mM ethylenediaminetetracetic acid were adjusted to 1.4 M sodium chloride and to 20 mM ethylenediaminetetracetic acid in extraction buffer. Despite these adjustments and multiple washing steps, the extracted RNA solution still contained contaminants with ratio 260/230 nm lower than 1.5 and thus had to be purified with glycogen (RNA grade, Thermo Fisher Scientific). Purification with glycogen was performed according to manufactures protocol with changes. Incubation step was performed in room temperature not at −20°C. Thanks to this adjustment, the extracted RNA precipitated but the contaminations remained desolved in the working solution. After precipitation, washing step was performed at least 2×. RNA extraction was performed in technical duplicates, so 28 RNA extractions was used for next step. Library preparation with polyA selection was performed with Dynabeads® mRNA Purification Kit and NEBNext Ultra Directional RNA Library Prep Kit for Illumina according to manufacture protocol. Indexed cDNA libraries were sequenced in one run following manufacturer’s instructions to generate paired-end, 150-base reads for reference transcriptomes and in two interdependent runs for single-end, 100bp (base pair) reads. Sequencing runs were performed on a HiSeq 4000 (Illumina). The datasets generated and analyzed during the current study are available in the NCBI database repository TSA: GJBZ00000000, GJBY00000000, GJBR00000000, GJBQ00000000, GJBP00000000, GJBO00000000, GJBN00000000.

### Transcriptomes assembly

Transcriptome assembly was performed based on recommendations for de novo assembly of non-model species (46). The quality of the reads and the effects of trimming were assessed using FastQC v0.11.8 (47). Raw reads were trimmed using Trimmomatic v0.39 (48) to remove low quality bases (phred score > 30), adapter sequences, and other sequencing artifacts. After this filtering step, biological replicates were merged to one for each individual, reads were then combined between libraries of the same species. In next step, we filtered out reads based on overall score (phred score > 25) per read, other parameters run on default settings. The erroneous k-mers from trimmed and filtered Illumina paired-end reads were corrected using Rcorrector v1.0.4. (49).

The corrected paired reads were assembled using Trinity v2.8.4 (50) (minimal contig_length: 300; group_pairs distance: 250; minimal kmer_cov: 2) separately for each species (2 samples together). First, the overlapping reads are combined into contigs with a minimum length of 300bp and at least 2 reads to be assembled. Subsequently, these contigs were reassembled into longer sequences of so-called unigenes. Because multiple samples from the same species were sequenced, the longest sequences were grouped into clusters based on the detection of splice variants and abundance. But because splice variants may be incorrectly classified as paralogs (as they can be assembled into different contigs but with the same component name in the assemblies) we retained only the longest isoform for each transcript, thus reducing this assembly error and making sure that each gene was represented by only one transcript. Evaluation of the structure of the generated assemblies was done with the QUAST software (51).

### Functional annotation and gene ontology analysis

Assembled non-redundant unigenes were functionally assessed for transcriptome completeness using Benchmarking Universal Single-Copy Orthologs (BUSCO (52)), which uses the entire embryophyte dataset, which represents the evolutionary informed near-universal single copy orthologs from OrthoDB v9. For quality assessment, we ran RSEM-EVAL from DETONATE software package (53). Based on RSEM-EVAL *contig impact score*, we removed all contigs with negative score from the assemblies and reran in RSEM-EVAL. Annotation was performed using the Trinotate pipeline (https://trinotate.github.io/). The coding regions in the transcripts and their most likely candidates for the peptides with the longest open reading frames (ORFs) were predicted using Transdecoder v.2.0.1 (54). Only the putative ORFs that were at least 100 amino acids in length were retained. Subsequently, sequence homology between predicted protein sequences and the NCBI non-redundant protein database (Nr) (ftp://ftp.ncbi.nih.gov/blast/db/) and Swissprot-Uniprot database was searched. The functional annotation was achieved using Hmmer v.3.1b1, a protein domain identification (PFAM) software (55) and Rnammer v.1.2 to predict ribosomal RNA (56). Sequences with a match in either NCBI non-redundant protein database (Nr) or Swissprot-Uniprot database were annotated with gene ontology (GO) terms using the PANTHER (protein annotation through evolutionary relationship) classification tool (57).

### Gene families, selection and candidate genes

We used the program OrthoFinder v1.1.2 (58) to identify orthogroups and orthologs of proteins among our 7 assemblies using Basic Local Alignment Search Tool (BLAST) all-v-all (self and reciprocal BLASTs simultaneously) and to derive rooted gene trees for all orthogroups and to identify all of the gene duplication. OrthoFinder analysis was conducted for all pair-wise comparisons, for all 7 species assemblies in order to identify putative orthologs within the current datasets. We used the outputs from OrthoFinder to determine the number of overlapping (shared across species) transcripts across the 7 assemblies.

The protein sequences associated with the ortholog group members were aligned using MAFFT v7.407 (59) and the corresponding coding sequence was matched and retrieved to ortholog alignment with PAL2NAL v14.1 (60). Subsequently the alignment was used to construct an maximum likelihood phylogenetic tree using RAxML (Randomized Axelerated Maximum Likelihood) v8.2.12 (61). The algorithm selected for this study conducted a rapid Bootstrap analysis and searched for the best-scoring maximum likelihood tree in one single program run. The selected substitution model was generalized time reversible (GTR) GAMMA. To maximum likelihood to statistically estimate dN and dS was used the CODEML program within the Phylogenetic Analysis by Maximum Likelihood (PAML) package (62). Which uses the observed changes present in the codon alignments from PAL2NAL, given the phylogenetic tree constructed by RAxML. CODEML calculates the likelihood of the observed changes resulting from two models of evolution, only one of which allows for the possibility of detecting positive selection (dN/dS > 1). GO terms were assigned for positively selected orthogroups as described above.

## Results

### De novo transcriptomes and assembly

RNA-seq libraries representing leaf transcriptomes yielded between 7 981 717 and 15 552 519 paired end reads per individual, for 2 individuals per species cultivated in contrasting temperature regimes. Between 1 and 26% of raw reads were removed due to quality filtering (quality score < 30). For each of the seven species, high quality RNA-seq datasets, which contained between 53 742 722 and 78 519 184 pair end reads were used for the assembly (Table 1).

The de novo assemblies (using Trinity) ranged between 92 035 and 226 081 contigs for the seven species (Table 2). More than 80% of the input reads mapped back to their assembled transcriptome, which is an indication of a good quality assembly. Similarly also RSEM-EVAL results (Table 2) showed proper assembly.

**Table 2.** Detonate RSEM-EVAL evaluation of results for each species

| Species | Number of contigs | RSEM-EVAL score | Number of contigs with no read aligned to | Number of alignable reads |
| --- | --- | --- | --- | --- |
| *I. balsamina* | 190 736 | -5 610 093 097.24 | 15 036 | 16 115 961 |
| *I. racemosa* | 194 470 | -6 480 726 545.38 | 20 400 | 16 980 323 |
| *I. bicornuta* | 226 081 | -8 102 571 944.57 | 25 347 | 17 092 225 |
| *I. scullyi* | 136 472 | -8 759 635 212.49 | 12 326 | 19 586 591 |
| *I. scabrida* | 94 707 | -7 102 916 062.01 | 11 122 | 12 472 536 |
| *I. falcifer* | 161 825 | -7 492 222 641.16 | 18 948 | 18 476 547 |
| *I. devendrae* | 141 792 | -7 866 313 302.92 | 18 852 | 12 712 480 |

### Functional annotation, gene ontology analysis and quality assessment

To perform functional annotation, the filtered assembly was submitted to the Trinotate pipeline and we predicted open reading frames (ORFs) with Transdecoder. Between 42.9 and 68.7% transcripts of *Impatiens* species were identified, resulting into 52.3-72.4% predicted protein coding genes (Fig 1). A protein gene set completeness assessment (BUSCO pipeline) showed that majority of *Impatiens* core genes had been successfully recovered in assemblies (Table 3). Only between 13.6 and 22 % of the *Impatiens* single-copy orthologs were classified as missing, suggesting good coverage and high quality of the assembly of the protein-coding transcriptomes.

**Fig.1.**
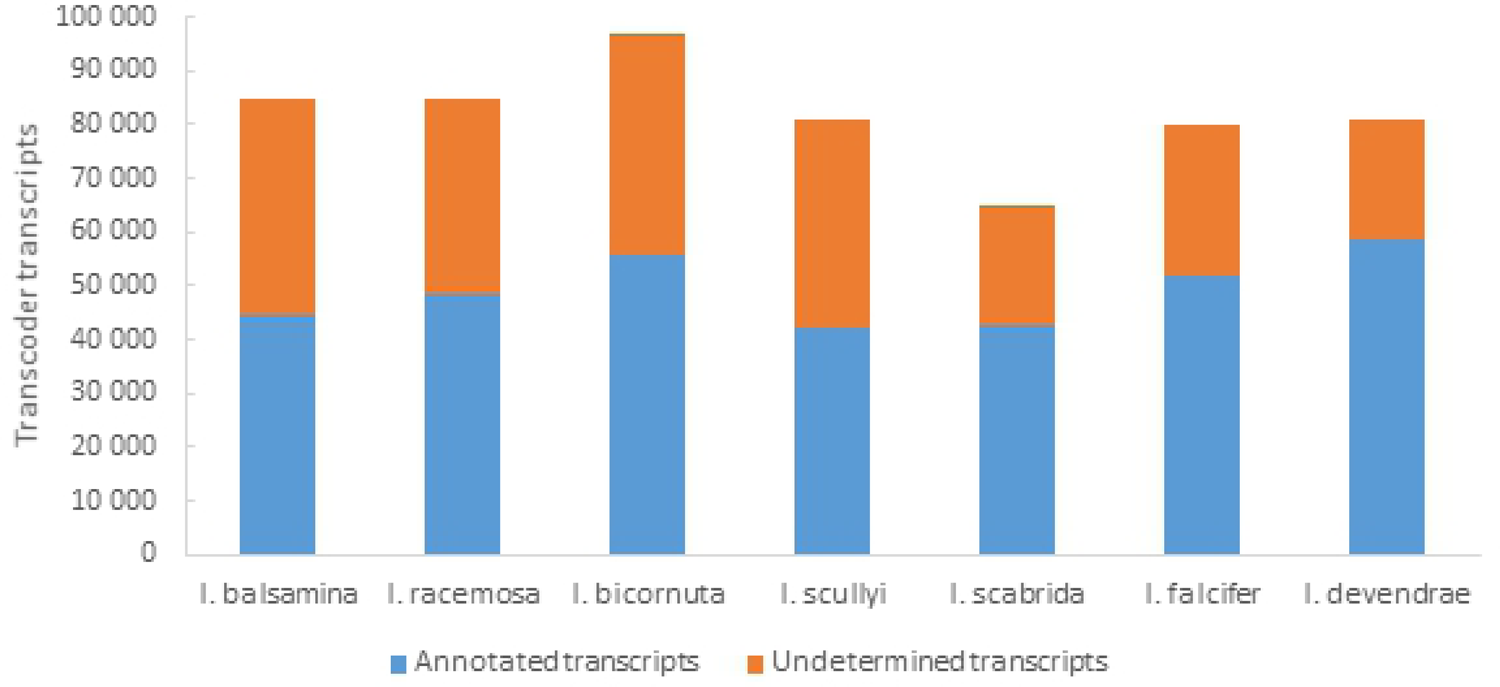
Annotation summary. Quantity of assembled transcripts with identified ORF by Transdecoder (Trinotate pipeline). Percentage values are showing proportion of annotated and undetermined transcripts for each species.

**Table 1.** BUSCO- an analysis of assembly completeness.

|  | *I. balsamina* | *I. racemose* | *I. bicornuta* | *I. scullyi* | *I. scabrida* | *I. falcifer* | *I. devendrae* |
| --- | --- | --- | --- | --- | --- | --- | --- |
| Complete genes (%) | 73.5 | 76.7 | 79.1 | 77.0 | 64.7 | 80.0 | 72.4 |
| Single-copy genes (%) | 40.5 | 40.9 | 31.0 | 42.5 | 31.9 | 34.9 | 33.4 |
| Duplicated genes (%) | 33.0 | 35.8 | 48.1 | 34.5 | 32.8 | 45.1 | 39.0 |
| Fragmented genes (%) | 10.3 | 7.2 | 6.2 | 8.1 | 13.3 | 6.0 | 9.7 |
| Missing genes (%) | 16.2 | 16.1 | 14.7 | 14.9 | 22.0 | 14.0 | 18.0 |

Annotated genes were assigned to GO terms, with the highest proportions of mapped GO terms for the current *Impatiens* transcriptomes related to binding (∼28.3%) and catalytic activity (∼50.4%) under “Molecular Function”, cellular (∼32%) and metabolic (24%) processes under “Biological Process”, and cell part (∼48.5%) and organelle (∼36.5%) under “Cellular Component” (Fig. 2 and Supplementary Table S2).

**Fig. 1.**
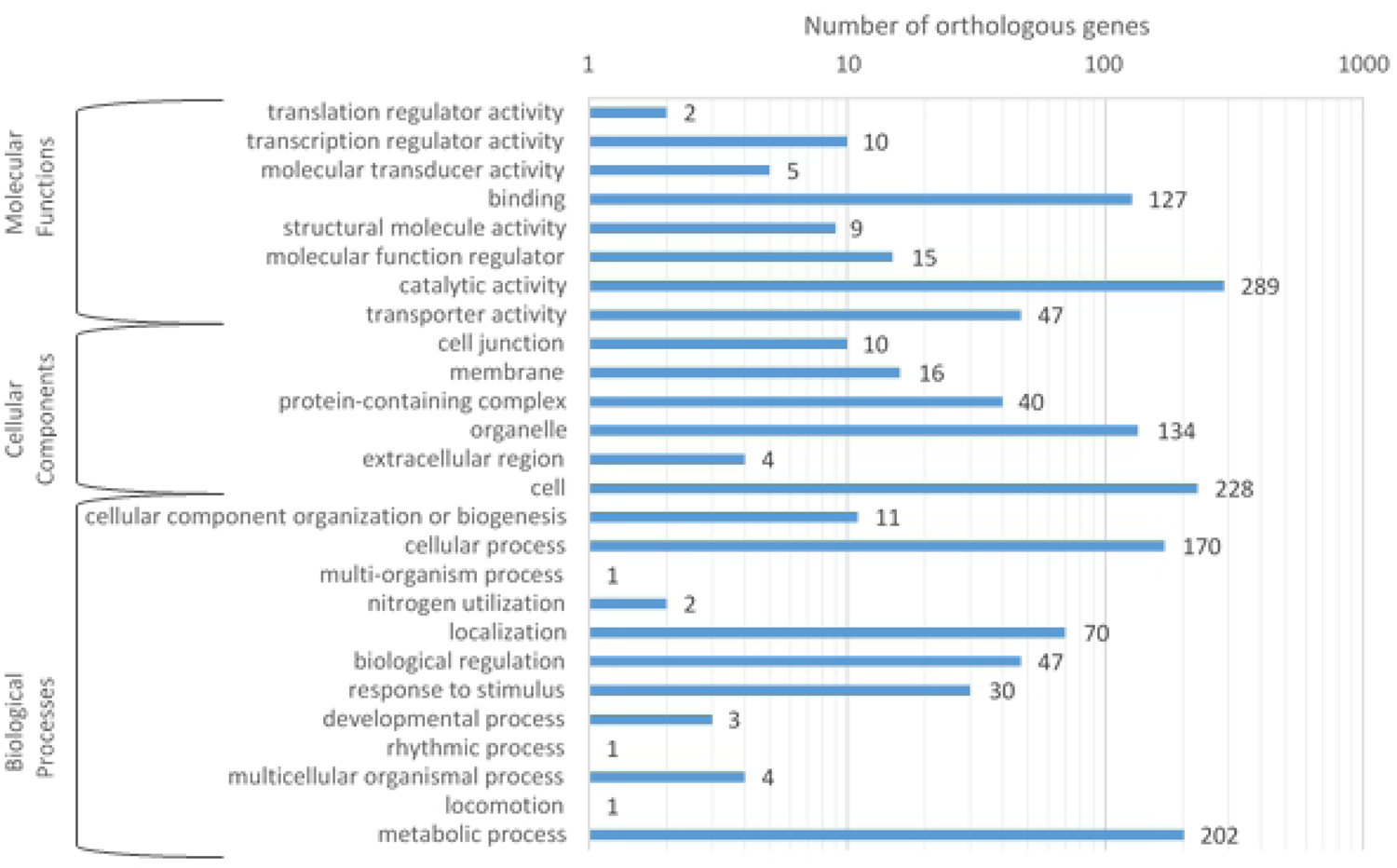
Histogram of GO terms of protein coding genes for each species.

### Gene families, selection and candidate genes

Total of 14 728 orthogroups (OrthoFinder) were found to contain *Impatiens* proteins, 77.5% of these also included one or more *Arabidopsis* proteins. A search for species specific orthology revealed a set of 236 unique groups that we consider as *Impatiens* specific. All species share 14 983 orthogroups, 5 246 orthogroups are missing in 1 species, 4 164 orthogroups are missing in 2 species, 6 803 orthogroups are missing in 3 species, 7 696 orthogroups were found in only 3 species, 11 518 orthogroups were found only in 2 species and 3 020 orthogroups were detected only in one species (Fig. 3, using OrthoFinder).

**Fig. 3.**
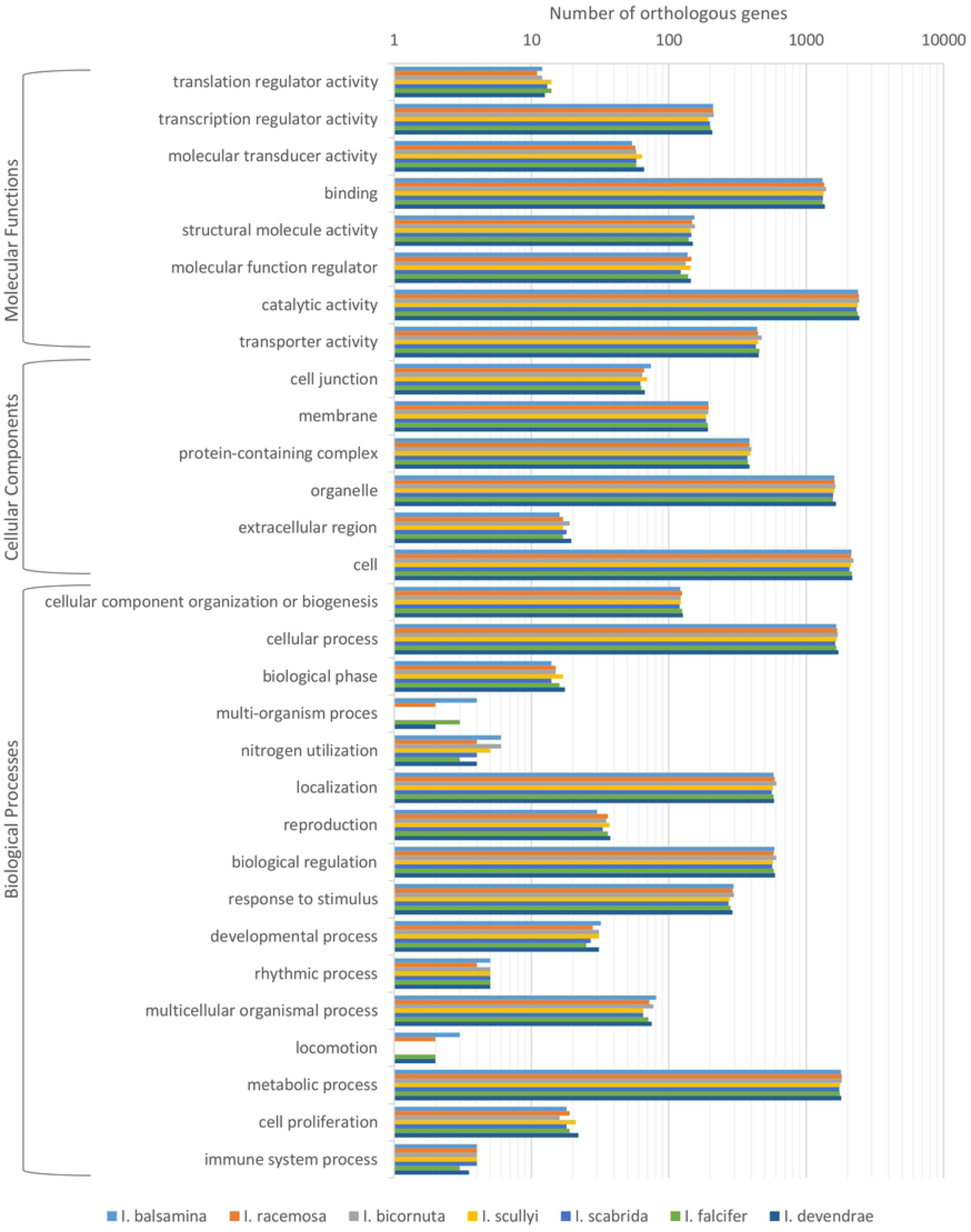
Orthogroups overlap between all seven species. Grey boxes in lines with species names indicate presence of the given orthologous group in this species.

For each species, the CODEML program identified between 2 536 and 3 009 orthogroups under positive selection (Supplementary Table S3). These positively selected groups were assigned to GO categories. The highest proportions of mapped GO terms were related to binding (∼28.2 %) and catalytic activity (∼51.5%) under “Molecular Function”, cellular (∼31.2 %) and metabolic (35,3%) processes under “Biological Process”, and cell part (∼49.5%) and organelle (∼35.5%) under “Cellular Component” (Fig. 4).

**Fig. 4.**
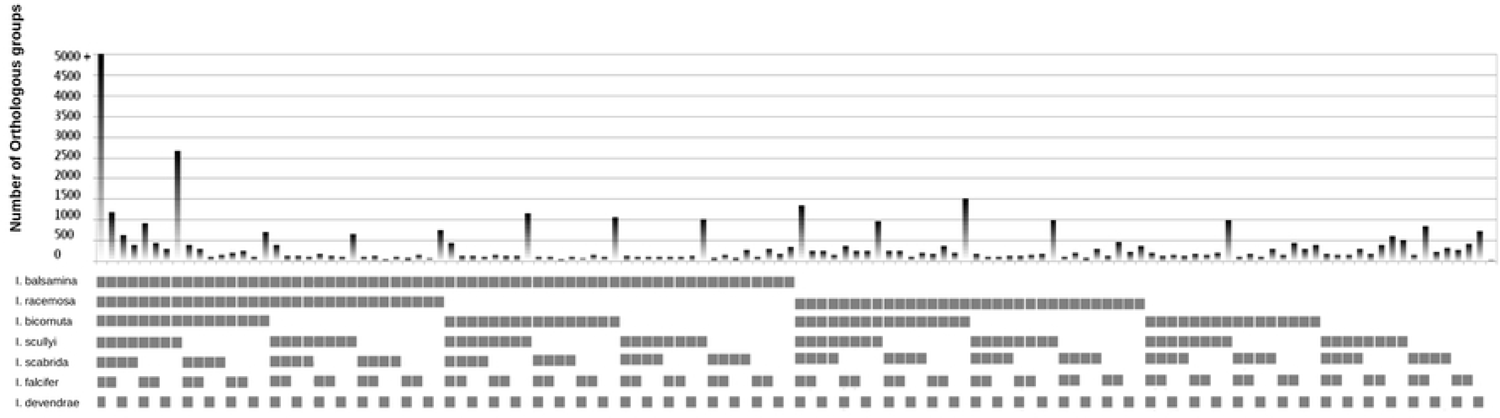
Histogram of GO terms of the clusters of orthologous groups under positive selection for each species.

Also 767 orthogroups under positive selection common for all investigated species were identified. The highest proportions of mapped GO terms for these orthogroups were related to binding (25.2%) and catalytic activity (57.3%) under “Molecular Function”, cellular (31.4%) and metabolic (37.3%) processes under “Biological Process”, and cell part (52.8%) and organelle (31%) under “Cellular Component” (Fig. 5 and Supplementary Table S4).

**Fig. 5.**
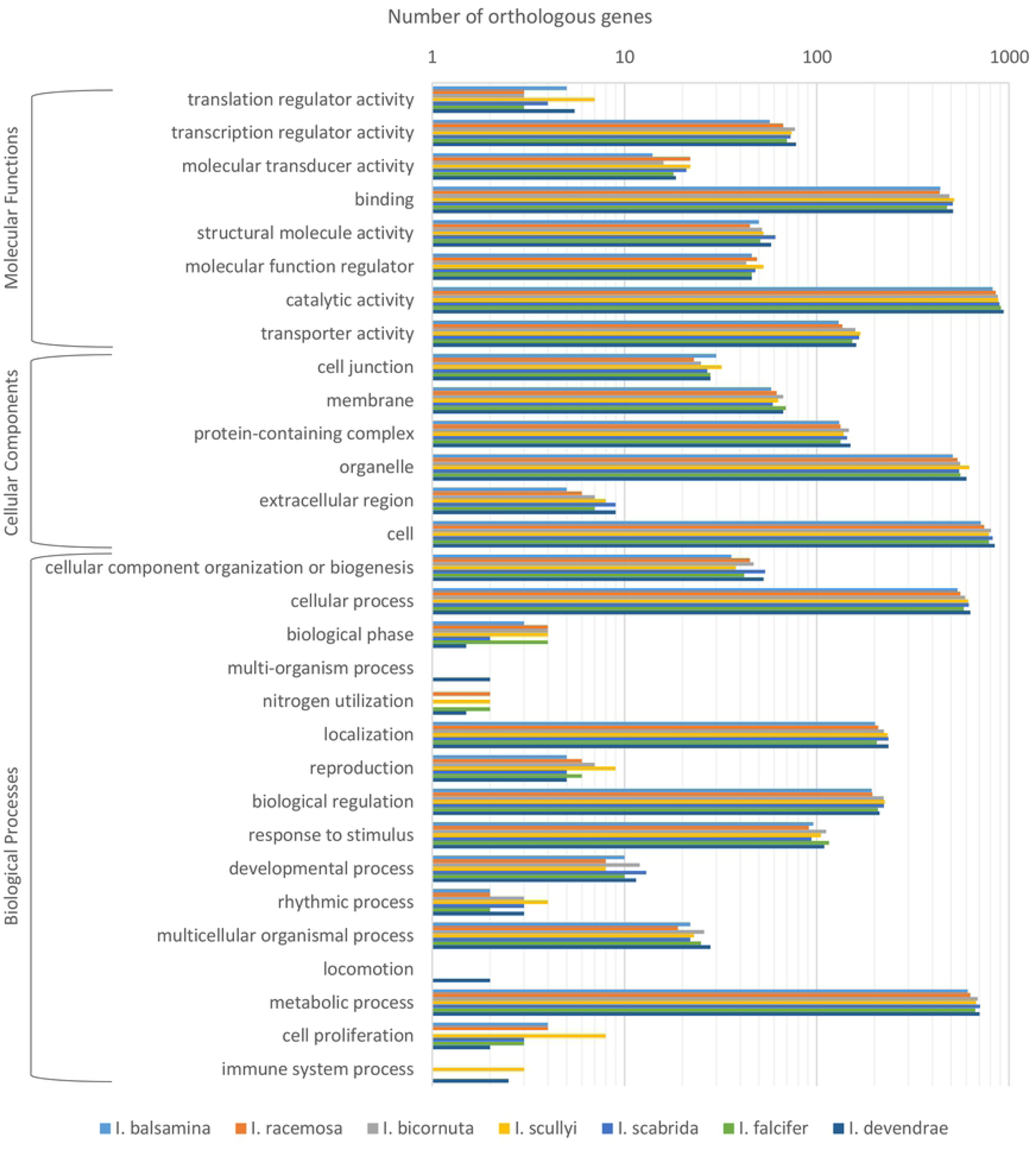
Histogram of GO terms of the clusters of orthologous groups under positive selection common for all species. Numbers represent number of identified orthologous genes for each category.

## Discussion

We performed RNA sequencing of 7 closely related species of genus *Impatiens* to assess the differences and similarities in the transcriptome within the group. The results indicate good quality of assembled transcriptomes with comparable annotations across the species. From identified orthologous gene families specific for *Impatiens* species, most abundant known motives were reverse transcriptase motive, gag-polypeptide of long terminal repeat (LTR) copia-type and Zinc knuckle motive. Six of the seven investigated species shared 77% of gene families. Despite *I. bicornuta* distinct selection profile, this species shared selection on zing finger protein structures and flowering regulation and stress response proteins with the other investigated species. This consistency may suggest that the group evolved via adaptive radiation.

Also similar transcriptome profile of *I. balsamina* compare to rest of the species was identified, despite this species belongs, based on phylogeny Yu et al. (43), to the G clade, rest of the species belong to clade B. This finding can be explained by study of Janssens et al. (23). They hypothesized that rapid radiation of this genus is caused by a periodicity of glacial cycles during the Pleistocene. That could have resulted in change of biotopes, with movement of vegetation belts. To inhabit suitable localities, *Impatiens* had to migrate along with the rainforest belts. We can expect, that climatic episodes isolated many different *Impatiens* populations for several thousands of years. This explanation leads us to possible scenario that species diversification is led by habitat requirements in given time in small isolated populations. At present, these populations do not suffer from long term climate pressure, so diversification and changes in stably transcribed genes can be slower, less pronounced, but still running.

Natural individuals are rarely studied, but Baker et al. (20) studied transcriptomes of four pine species. Some of the identified families are in common with our findings (F-box family, ubiquitin related proteins, stress response family). Li et al. (63) compared more differentiated species in *Rosaceae* (*Rosa, Malus, Prunus, Rubus,* and *Fragaria*) and they identified rapidly evolving genes responsible for DNA damage repair, respond to environment stimuli and post hybridization genome conflicts.

### Characterization of de novo sequenced transcriptomes and between species transcriptome profiles comparison

#### De novo transcriptome reconstruction

Despite necessary changes in RNA extraction protocol due to high concentration of secondary metabolites in the plant tissue, we were able to obtain good quality RNA extracts. Also high proportion (average 96%) of preserved reads after quality trimming, which is comparable with *I. balsamina* trimming (93%) in (40) indicates high quality of extracted RNA. Overall, RNA seq library construction with implemented changes was successful.

Comparison of the reads and subsequent assemblies showed many similarities in summary statistics among the sequenced transcriptomes. The biggest noticeable difference is in the size of the largest identified contigs. For three species, they are significantly smaller than in the other four. It can be caused by lower differences between isoforms, alternative splice forms, alleles, close paralogs, close homologs, and close homeologs compared to the other species. Six of the seven studied species have not been sequenced before. Only *I. balsamina* transcriptome has been recently sequenced (40). The transcriptome size we obtained for *I. balsamina* is similar to what has been previously published (84 635/ 91 873 transcripts).

*Impatiens bicornuta* has the biggest detected transcriptome from all the investigated species (226 081 contigs) and also the longest assembly (196 656 333 bp). The smallest transcriptome and shortest assembly were identified for *I. scabrida* (94 707 contigs and 95 826 671 bp). Number of published reference transcriptomes for this genus is very limited. Transcriptome of *I. walleriana* (64) was also previously sequenced and its transcriptome size (121 497 contigs) is comparable with our results. As all our species were cultivated in identical conditions, the differences in transcriptome size between species can be caused by different habitats of plant origin and history of each species which cause changes in gene expression due to gene expression regulation, epigenetic mechanisms, transposon activation, sequence changes, different combinations of regulatory factors and/or increased gene dosage. Nagano et al. (65) showed intra-annual fluctuation transcriptome size in *Arabidopsis halleri subsp. gemmifera*. They described from 2 879 to 7 185 seasonally oscillating genes. Gurung et al. (66) compared transcriptome size in a Himalaya plant (*Primula sikkimensis*) growing along an altitudinal gradient and carried out a transplant experiment within the altitudinal gradient. Total number of transcribed genes fluctuated between 38 423 and 48 674 depending on altitude of plant cultivation, which reflects species plasticity rather than adaptation.

The 86.7 % of contigs, which were correctly assembled corresponds to other assembled transcriptomes: 95.1% complete genes in (67), 91% in *Noccaea caerulescens* (68) and 94.4% in *Centaurium erythraea* (69). The highest (22%) proportion of missing and fragmented genes was identified for *I. scabrida.* This can be partially explained by mean length of the assembly, which is the highest (1021bp) for this species.

In terms of annotation, *I. devendrae* has the lowest percentage of undetermined transcripts (27.6%). Higher percentage of undetermined transcripts for the rest of the species is common for annotation without a reference genome (70). Higher number of unannotated transcripts can represent short or newly identified contigs which cannot be uniquely identified in public databases. We annotated more than 73 000 genes for each species, which is comparable with *I. walleriana* result (70 190). Our results are also comparable with other genera in *Ericales* order such as *Primula* (67 201 genes, (66)), *Phlox* (59 994 genes, (71)), *Saltugilia* (51 020-92 672 genes,(72)), *Camellia* (46 223-46 736, (73)) and *Vaccinium* (35 060 – 67 836 genes (74, 75). GO terms assignment for identified transcripts is surprisingly similar between\ species. We expected that species differentiation and origin (altitudinal gradient) of sequenced individuals will affect overall transcriptome profiles. Our results suggest that species within this genus transcribe similar sets of genes and potential differences can be pronounced in differential expression of these genes or their sequence. Similar results were published by Gurung et al. (66), who studied species plasticity by comparison of transcriptome size in *Primula sikkimensis* and they identified 21 167 transcripts and 109 and 85 genes which were differentially expressed between three altitudinal positions in which the plants were cultivated.

#### Orthologous gene families (Impatiens specific)

From identified orthologous gene families (14 728) for each species, we identified 334 gene families specific for *Impatiens* species. In these, 143 specific protein motives have been found. Fourteen of them were unknown, most abundant known motives were reverse transcriptase motive (7 times), gag-polypeptide of LTR copia-type (6) and Zinc knuckle motive (5). In group of 3 most common motives are integrase core domain, LTP family and RNA recognition motif. Majority of identified motives were found only once, so we will focus in this discussion on the most abundant hits only.

Reverse transcriptase, gag-polypeptide of LTR and integrase motive is expected in identified transcripts (76). These enzymes are generally connected with viruses and retrotransposons without specific function in plant cell and they are transcriptionally active. Lescot et al. (77) published that reverse transcriptase genes (RT) are more abundant in plankton metatrascriptome compare to a metagenome with distinct abundance patterns. They proposed that transcription of various RT-assisted elements could be involved in genome evolution or adaptive processes of plankton assemblage. Another possible source of transcripts was suggested by Gladyshev and Arkhipova (78) providing evidence of single copy reverse transcriptases. They found corresponding sequence in all major taxonomic groups including protists, fungi, animals, plants, and even bacteria evolving under strong selective pressure.

Their function is not experimentally confirmed. Gag polypeptide motive is in congruence with reverse transcription genes. They both are parts of retrotransposomes. These elements are studied in connection to environmental stress. Kalendar et al. (79) found correlation between LTR insertion in barley genome and ecogeographical distribution connected with BARE-1 promoter of abscisic acid-response elements typical for water stress-induced genes. Reverse transcriptase genes and LTRs opens interesting field for a molecular research to reveal their function and regulation in transcription level.

Zinc knuckle motive is studied from many perspectives. Loudet et al. (80) demonstrated that zinc knuckle protein negatively controls morning-specific growth in *Arabidopsis thaliana*.

Also, proteins with this motive were found as a component of alternative splicing machinery in response to osmotic and salinity stress (81, 82). Sequenced *Impatiens* species originated in different altitudes from Himalayas in Central and East Nepal. Osmotic or salinity stress is not expected environmental factor to shape transcription in these regions. More probably environmental factors as higher dosage of UV light and reactive oxygen species (83) will potentially change genome and/ or transcriptome response. Indeed, Zhao et al. (84) identified selection on zinc knuckle protein family as a part of genetic adaptation of *Lobelia aberdarica* and *L. telekii* transcriptomes to different altitudes in East African mountains as a response to DNA damage caused by volcanic eruptions, UV and frost damage. Zinc knuckle protein family (CX2CX3GHX4C) belongs to zinc finger superfamily (CCHC-type), so potential for functional diversity of this protein family is large.

Lipid transfer proteins (LTP family) and RNA recognition motives are involved in many processes, but two of them are common for both of them-pathogen defense and response to environmental stress (cold). Plants are continuously evolving complex regulatory networks to respond to pathogen threats (85) so specific *Impatiens* sequence changes are expected.

Similarly, adaptation to cold in locations of origin of the investigated *Impatiens* species (1 330-2 728 m above sea level) is probably strong ecological driver of genome/transcriptome changes. Specifically, lipid transfer proteins are involved in the intra- and extracellular transport of lipids (86). Hincha et al. (87) published that member of LTP family, cryoprotectin is involved as a protection of thylakoid membranes during a freeze-thaw cycle in cabbage.

Another member of LTP family (LTP 3) was also identified to be involved in response to freezing in model species *A. thaliana* (88). Multiple proteins with glycine-rich RNA binding proteins were found active during cold acclimation e.g. AtGRP7 (89), AtGR-RBP2 and AtGR-RBP4 (90).

### Gene families under positive selection and differences between species

Six of the seven investigated species shared 77% of gene families. Below, we on the 23% of protein families differing among species and thus likely under selection and on distinct *I. bicornuta* profile and their involvement in response to different environment stimuli. 50S ribosome-binding GTPase domain is not discussed because lack of information.

### Resistance

Resistance (R) proteins are involved in pathogen recognition and activation of immune responses. Most resistance proteins contain a core nucleotide-binding domain. This part is called NB-ARC domain and consists of three subdomains: NB, ARC1, and ARC2. The NB-ARC domain was identified under selection only in *I. balsamina* transcriptome. It is a functional ATPase domain, and its nucleotide-binding state regulates protein activity (91).

Recently, Ghelder et al. (92) reported fusion of an RPW8 (resistance to powdery mildew 8) domain to a NB-ARC domain in conifers as a part of response to drought. Similarly, Bailey et al. (93) performed phylogenetic analysis on plant immune receptors with NB-ARC domains originated in grasses. They described that plant immune receptors are able to recognize pathogen effectors through the acquisition of exogenous protein domains from other plant genes through DNA transposition and/or ectopic recombination.

*Arabidopsis* broad-spectrum mildew resistance protein RPW8 is coded by naturally polymorphic locus and also belong to R proteins. Although genes are polymorphic, protein structure is stable (94). Zhong and Cheng (95) investigated RPW8 genes in 35 plant genomes and they found evidence for series of genetic events such as domain fissions, fusions, and duplications. Species-specific duplication events and tandemly duplicated clusters are processes responsible for species specific expansion.

### Response to heavy metal stress

In *I. balsamina* heavy-metal-associated domain was found under selection. As name of the domain suggests, it is part of heavy-metal-associated proteins and they are involved in heavy metal detoxification. Li et al. (96) identified six clades in *A. thaliana* and *O. sativa* based on specificity of heavy metal-associated domains. They also identified tandem and segmental gene duplication and *cis*-acting elements on close proximity of promoter sequences. This finding suggests regulation by multiple transcription factors. Similar results were also published on *Populus trichocarpa* (97).

### Protein families with zinc finger motive

B-box family was found under selection in two species: *I. scullyi* and *I. devendrae*. B-box family contains 32 zinc finger transcription factors which are involved in regulation of hormone signaling pathways and through them they regulate wide spectrum of processes such as drought-induced flowering pathway, germination, seedling photomorphogenesis, hypocotyl elongation in seedlings, induction of flowering, regulation of branching and shade avoidance response and many more. Despite conserved domain topology, this family is susceptible to environmental changes through variation in promotor region. In this region several different cis elements can be incorporated e.g. ABRE (ABA - Responsive element), ERE (EthyleneResponsive Element), CGTCA-motif and TGACG-motif (related to methyl-jasmonite stress responsiveness). Rosas et al. (98) identified natural variation in cis regulatory sequence of CONSTANS transcription factor which regulates flowering time in *Arabidopsis*. They also showed recent evolution of mutation and its spread to high frequency in *Arabidopsis* natural accessions.

Zn-finger in Ran binding protein family was detected in *I. balsamina, I. scullyi* and *I. scabrida*. This protein family is very poorly studied. Part of proteins belonging to this family is still uncharacterized. One of the characterized protein subfamilies in this family is organellar zinc finger subfamily. These proteins are located in plastid and mitochondria. Their function is not fully defined yet, but they are involved in chloroplast RNA editing (99).

Another example of Zn-finger motive in binding protein is histone deacetylase 15, which are involved in the repression of chlorophyll synthesis in the dark (100). Also TATA-Binding Protein-Associated Factor 15/15b belong to this group and is known to suppress flowering during vernalization (101). More described in molecular functions is Ariadne subfamily in Zn-finger family, a group of E3-type ubiquitin ligases, involved in last step of protein degradation (102).

### DNA repair and protein translation

RecA proteins were identified under selection in all studied species except *I. scullyi*. Recombinases are responsible for homologous recombination and maintenance of genome integrity. Recombinase RecA is crucial for DNA repair. RecA bind to ssDNA break and scan for a homologous template dsDNA (103). Miller-Messmer et al. (104) studied DNA recombination events in mitochondria. They found that RecA-dependent repair has a dual effect on the mtDNA: maintaining the integrity of the mitochondrial genome and also preferring the amplification of genome organization that could help in the adaptation to environmental conditions.

CAAD domain of cyanobacterial aminoacyl-tRNA synthetase was found under selection in *I. balsamina, I. racemosa* and *I. falcifer*: Aminoacyl-transfer RNA (tRNA) synthetases are key molecules in translation and act early in protein synthesis by mediating the attachment of amino acids to tRNA molecules. In plants, multiple versions of the protein are coded because protein synthesis is running in three subcellular compartments (cytosol, mitochondria, and chloroplasts). Aminoacyl-transfer RNA (tRNA) synthetases are acquired in *Arabidopsis thaliana* genome through horizontal gene transfer event from bacteria. These genes in plant evolution were under selection to maintain original function, while additional copies diverged (105).

### Regulation of flowering

CCT (chaperonin containing TCP-1) gene family was found under selection in *I. balsamina, I. racemosa, I. scabrida* and *I. falcifer*. Most CCT genes regulate flowering and they are studied mainly on rice for agricultural reasons (106) but studies on *Aegilops tauschii* proposed that CCT genes play important role in adaption of *A. tauschii* to the photoperiods of different regions. Zheng et al. (107) identified rapid evolution of CCT genes. More gene variations are available for adaption to environmental changes, so positive selection has kept more convenient variations.

### Nitrate detection and metabolism

Fip1 motive was found under selection in *I. balsamina, I. racemosa* and *I. falcifer*. Fip1 gene plays important role in nitrate signaling through expression regulates of kinesis CIPK8 and CIPK2 in *Arabidopsis* (108). Nitrate content in environment is very important indicator and regulatory networks consist of approximately 300 genes differently regulated under variety of conditions (109). These findings showed that nitrate signaling is under environmental pressure and adaptability to nitrate uptake, transport, assimilation and metabolism is crucial.

### Cell wall and membrane polymers

N-acetylglucosamine transporters were identified under selection in *I. balsamina, I. scullyi* and *I. falcifer*. The biosynthesis of glycans, glycoproteins and glycolipids requires glycosyltransferases localized in the Golgi apparatus and endoplasmic reticulum (110). They were identified as overexpressed in rice after salt stress exposure (111) and are also required for an arbuscular mycorrhizal presymbiotic fungal reprogramming (112).

### Families with pleitropic effects

Ras superfamily domain and GTPase domains were identified under selection in all investigated species. Ras superfamily consists of Rab, Ras, Arf, Rho and Ran families in yeast and animals, but the Ras family has not been found in plants. This superfamily is one of the most important gene families involved in signal transduction, vesicle trafficking, signaling, cytoskeleton rearrangements, nuclear transport, cell growth, plant defense in interaction with microorganisms (113). Their important role is also in adaptation to oxygen deprivation through levels of H_2_O_2_. H_2_O_2_ acts as a second messenger for either activate expression of genes responsible for tolerance against oxygen deprivation (e.g. the gene encoding alcohol dehydrogenase) or activate the generation of intracellular Ca^2+^ signals responsible for cell growth (114).

Specific selection profile of *I. bicornuta*

### Stress response

AP180 N-terminal homology (ANTH) domain is studied just recently and their functions are not fully understood yet. *A. thaliana* genome contains 18 ANTH proteins which showed their possible diversification in comparison with fewer ANTH proteins in metazoan and fungal genomes (115). Nguyen et al. (116) showed that mutant in ANTH protein, ateca4 showed higher resistance to osmotic stress and more sensitivity to exogenous abscisic acid molecules.

Hsp20 domain belongs to small heat shock proteins that are regulated by stress and work as molecular chaperons to protect proteins from stress related damage (117). Some of them evolved only recently, others are ancient (118). This region is conserved in C-terminal, but in 2 subfamilies β6 sheet is missing in plants and some animals. This motive is necessary for dimerization and oligomerization (119), but investigated proteins are able to create dimers anyway (120). On the other hand, N-terminal domain is very variable. Despite unstable structure, this region is functionally important because of substrate binding specificity (121). It is possible to conclude that these proteins have capacity for diversification.

Stress responsive A/B Barrel Domain forms dimers and its function is not fully understood. Shaik and Ramakrishna (122) found this domain/protein co-expressed under drought and bacterial stresses in *A. thaliana* and *Oryza sativa*. Protein with this domain was also identified downregulated in salt stressed *Abelmoschus esculentus* L. seedlings (123).

Uroporphyrinogen decarboxylase (URO-D) is involved in chlorophyll biosynthesis (UniProtKB - Q93ZB6). Mock et al. (124) identified this enzyme as a part of plant defense respond to stress caused by reactive oxygen species in *Nicotinia tabacum*. Vanhove et al. (125) also identified this protein significantly abundant as response to drought stress in banana. Zhu et al. (126) found increased abundance in this protein in waterlogged *Vitis vinifera*.

### Cellular processes

Components of kinetochore, centromere proteins A are proteins under positive selection in plants. They contain DNA binding motive. Centromeres are built from repetitive satellite sequences that are rapidly evolving, so proteins with binding specificity must evolve accordingly (127). E-box proteins control protein ubiquitination by ubiquitin 26S-proteasome system. They are involved in wide spectrum of processes including secondary metabolite regulation, response to stresses, phytohormone signaling, developmental processes and miRNA biogenesis (128). Schumann et al. (129) did phylogeny analysis of largest F-box subfamily because in non-plant eukaryotes genomes only one copy is present compared to *A. thaliana* with 103 copies in genome. They identified signatures of positive selection and part of the diverse protein domain which is lineage specific. These findings showed adaptation potential of this family.

Maintenance of mitochondrial morphology protein 1 is very poorly studied. This protein is part of protein complex, which physically connect endoplasmatic reticulum and mitochondria. Wang et al. (130) showed upregulation of this gene in *Aspergillus nidulans* under salinity stress.

RimM (ribosome maturation factor) N-terminal domain belongs to RimM protein, which is associated with 30S ribosomal subunit. This indicates its function in translation initialization or subunit maturation (131). It is been found in some genomes (*Plasmodium falciparum* and *P. yoelii*, *Anopheles gambiae*, *A. thaliana* (132)) but very little is known about it.

### Protein families with zinc finger motive

FAR1 (FAR-RED IMPAIRED RESPONSE1) DNA-binding domain is an N-terminal C2H2 zinc-finger domain. It is part of well characterized transposase-derived transcription factor FAR1. This transcription factor is involved in many processes e.g. light signal transduction, chloroplast division, chlorophyll, myo-inositol and starch biosynthesis, circadial clock regulation and drought and low phosphate response (133). Salojärvi et al. (134) showed that this gene family can be involved in adaptation to environment in *Betula pendula*.

Zinc-finger of the FCS-type (phenyl alanine and serine residues associated with the third cysteine) form one of the largest transcription factor families in plants. They are categorized into subfamilies based on the order of the Cys and His residues in their secondary structures (135). Such a large family is involved in many stress related pathways. They have a role in salt, cold, osmotic, drought, oxidative and high-light stress. Specifically, Jamsheer et al (136) showed that downregulation of FCS-LIKE ZINC FINGER 6 and 10 is part of osmotic stress responses in *A. thaliana*. Emerson and Thomas (137) published analysis of the gene family across the tree of life and identified positive selection acting on this family and specifically selection to change DNA-binding specificity of transcription factors.

### Family with pleitropic effects

Homeobox domain is present in 14 classes of transcription factors in plants. They are involved in wide spectrum of processes (signal transduction networks involved in response to abscisic acid, auxin and drought, general growth regulation, in shoot and floral meristem development, stem-cell specification and proliferation, flowering time regulation directly repress gibberellin and lignin biosynthetic gene expression and in the adaptation to drought and disease resistance to fungal pathogens and many more (138). More recently Khan et al. (139) investigated involvement of homeobox genes in *Brassica rapa* under various stress conditions. They identified active purifying selection on this gene family and dynamic variations in differential expression combined with their responses against multiple stresses.

### Energy metabolism

Oxidoreductase FAD-binding domain is part of flavoprotein oxidoreductases molecule. They act in oxidative pathways and are able to oxidize NADH/ FADH_2_. These molecules are well studied in different organisms but recently Trisolini et al. (140) compared sequences and crystal structures across selected taxonomic groups (*Bacillales*, *Enterobacteriales*, *Rhodospirillales*, *Rhodobacterales*, *Thermales*,*Rhizobia, Nematoda*, *Mammalia*, *Arthropoda*, *Anthozoa*, *Fungi*, *Plants and H. sapiens*, *S. cerevisiae*, *C. thermarum*). They showed that these proteins have different coding sequences, but despite that, protein structure and shapes are similar and functional in oxidation processes.

### Flowering regulation

SNW domain in SKIP protein regulate pre-mRNA splicing and also was identified as a transcription factor of gene transcription activator in Paf1 complex, which is involved in transcription of FLOWERING LOCUS C and flowering in *A. thaliana* (10). It is involved in alternative splicing regulation of clock and salt tolerance-related genes in plants (141).

## Conclusions

We performed RNA sequencing of 7 closely related species of genus *Impatiens*. Most abundant motives were reverse transcriptase motive, gag-polypeptide of LTR copia-type and Zinc knuckle motive from orthologous, *Impatiens* specific gene families. Six of the seven investigated species shared 77% of gene families under selection. We also described distinct *I. bicornuta* selection profile, but this species shared selection on zing finger protein structures and flowering regulation and stress response proteins with the other investigated species. The reason for the strong differentiation of this species, however, remains unclear, but shared selection may indicate evolution via adaptive radiation in *Impatiens* species.

It is possible to conclude that genes involved in response to environmental stimuli is a major driver for selection in *Impatiens* species. More natural populations and species have to be sequenced to reveal conserved biological principles and distinguish lineage and locality adaptations.

## Acknowledgements

We thank to P. Vácha (Seqme) for excellent sequencing help, M. Rokaya for collection of the seeds used for the study and Z. Líblová for cultivation of the plants.

## Supporting Information file

Table S1.xls

List of species and location of their collection

Table S2.xls

Identified GO terms for each species.

Table S3.xls

Identified orthologues under selection for each species.

Table S4.xls

Identified orthologues under selection common for all species.

